# Effects of lipid membranes on RNA catalytic activity and stability

**DOI:** 10.1101/2024.08.31.610601

**Authors:** Tomasz Czerniak, James P. Saenz

**Affiliations:** Technische Universität Dresden, B CUBE Center for Molecular Bioengineering, 01307 Dresden, Germany; Technische Universität Dresden, Faculty of Medicine, Dresden 01307, Germany

## Abstract

RNA plays crucial roles in cellular organization and metabolism, and modulating its activity is essential for maintaining cellular functions. RNA activity, involving both catalytic (ribozymes) and translation processes, is controlled via myriad mechanisms involving different binding partners such as proteins and smaller polar solutes. We previously reported that lipid membranes can directly interact with the artificial R3C ribozyme changing its activity, however the effect of lipids on naturally occurring ribozymes remains unknown. Here, we report that both catalytic activity as well as RNA integrity can be controlled by the presence of different lipid membranes. Gel-phase lipid membranes decreased the activity of hepatitis delta virus (HDV) ribozyme and increased the activity of a hammerhead (HH) ribozyme. The presence of lipid liquid membrane surfaces triggered RNA degradation with greater degradation occurring in the single-stranded regions of RNA. The interplay between RNA activity and stability in the presence of different lipid membranes introduces multiple possibilities, where different combinations of ribozyme and lipid membrane composition could produce different effects on activity. Taken together, these observations support the hypothesis that the activity of both natural and artificial RNAs can be modulated by lipid membranes which, in turn, provides a foundation for the development of novel riboswitch-like molecules, and lipid membrane-based RNA-biosensors.

## Introduction

Ribozymes are catalytically active RNA molecules which play crucial roles in cellular as well as viral metabolism^1–4^. The activity of ribozymes is strictly dependent on correct RNA folding, and on the presence of interaction partners such as divalent cations. For example, hepatitis delta virus (HDV), a pathogen involved in liver cirrhosis development, contains the crucial ribozyme sequence involved in HDV replication in which activity is strictly dependent on a characteristic pseudoknot folding^5,6^ and the presence of divalent cations. Similarly, another class of ribozyme, hammerhead (HH), involved, for example, in viroid and retrozyme replication, also requires specific structures in order to be catalytically active, however it does not necessarily require divalent cations^7,8^. Ribosome and RNAseP activity are modified by interaction with proteins^9–11^. These phenotypes offer a range of possibilities in which different natural ribozymes can be controlled in different ways using relatively simple changes in the ribozyme microenvironment.

Most ribozymes are capable of maintaining their full catalytic activity without other macromolecules^1,12,13^. It was discovered, however, that RNA activity can be changed through the interaction with small solutes such as ATP, theophylline, and cyclic di-guanosine monophosphate in the special RNA classes such as aptazymes and riboswitches^14–17^. Moreover, recent research has shown that ribozyme activity can be also controlled through interactions with much larger binding partners such as single-component lipid membranes^18^. It is, however, unclear how different lipid membranes, for example varying in lipid composition or membrane fluidity, can further tune the activity of ribozymes.

Lipids, due to their amphiphilic character, spontaneously assemble as lipid bilayers, which form lipid vesicles (liposomes). Liposomes can act as a pseudo two-dimensional platform (scaffold) for binding different molecules such as metal ions, proteins, and nucleic acids. Lipid membranes can differ in their composition (e.g. phospholipids, sterols) which can further modify the features of the bilayer (e.g. fluidity, viscosity, diffusion). For example, a lipid membrane composed of lipids above their melting temperature (Tm) form a liquid disordered phase (L_d_) in which lipids can freely diffuse laterally within the lipid membrane matrix, whereas lipid membranes composed of lipids below their Tm form a gel membrane (s_o_) in which the movement of lipid molecules is impaired. Introduction of cholesterol molecules can further modify lipid membrane fluidity, in some cases creating so-called liquid ordered phase (L_o_) which is less fluid than L_d_ and more fluid than gel membranes^19^.

RNA-lipid interactions have been reported to involve electrostatic forces^20–22^, while hydrophobic interactions, such as RNA penetrating the lipid membrane surface, have also been proposed^23–27^. RNA-lipid interplay can result in functional outcomes and regulatory mechanisms that influence RNA phenotype and activity. For instance, the interaction, which can be tuned both by the RNA structure and sequence, could stabilize certain RNA conformations, which could lead to changes in ribozyme activity^18,25,26,28^. Thus, it is potentially possible to use lipids to control the activity of ribozymes with different reaction mechanisms, sequences, and structures.

The effect of lipid membranes on artificial R3C ligase^29^ was previously demonstrated, showing that changes in the sequence of R3C ligase substrate significantly changed characteristics of the ribozyme activity. Namely, the presence of lipid gel membranes improved both the interaction between R3C ligase and its short RNA substrate via sequence-dependent RNA-lipid binding^18^. It is, however, unclear how the presence of lipid membranes can influence the non-artificial ribozyme activity directly by the binding of the ribozyme molecules.

Here, we show how different ribozymes can be affected by interactions with lipid gel membranes. The activities of HH and HDV ribozymes were influenced differently. The HDV ribozyme showed inhibition, while the HH ribozyme exhibited an increase in reaction rate. We further report lipid-membrane dependent RNA-degradation which depends on membrane fluidity. Taken together, both activity changes (R3C, HH, HDV) as well as lipid-triggered degradation create a new level of control which might be employed in synthetic as well as molecular biology, and help to understand riboregulatory processes.

## Methods

### Materials

HEPES, magnesium chloride (MgCl_2_), calcium chloride (CaCl_2_), ammonium chloride (NH_4_Cl), crystal violet, Orange G, Tris, boric acid, ethylenediaminetetraacetic acid (EDTA), sodium acetate (NaOAc) were purchased in Carl Roth. Lead acetate (Pb[OAc]_2_), choline chloride, glycerophosphocholine (GPC), cholesterol (Chol) were purchased in Sigma Aldrich.

1,2-dioleoyl-sn-glycero-3-phosphocholine (DOPC),

1,2-dipalmitoyl-sn-glycero-3-phosphocholine (DPPC),

1,2-diarachidoyl-sn-glycero-3-phosphocholine (20:0 PC),

1,2-diphytanoyl-sn-glycero-3-phosphocholine (DPhPC),

2-((2,3-bis(oleoyloxy)propyl)dimethylammonio)ethyl ethyl phosphate (DOCPe),

1,2-dipropionyl-sn-glycero-3-phosphocholine (03:0 PC),

1,2-dioleoyl-3-trimethylammonium-propane (DOTAP) were purchased in Avanti Polar Lipids.

SybrGold was purchased in Thermo Fisher. All buffers were prepared in MilliQ water (Merck). All chemicals were used without further purification.

### RNA synthesis and purification

HDV and HH constructs were obtained using a standard *in vitro* T7 transcription (**Suppl. Table 1**), overnight incubation 37 °C. RNA was separated using denaturing polyacrylamide gel electrophoresis (PAGE), post-stained with 1xSybrGold, visualised with the blue light, and excised from the gel using the razor blade. Gel slices were then crush-and-soaked in the Tris-borate-EDTA buffer, 90 °C, for 1 hour. Obtained RNA was ethanol precipitated (0.3 M NaOAc), washed with 70% ethanol and resuspended in MilliQ water. Concentration and integrity of RNA were estimated using absorbance readout and PAGE, respectively. A 6-carboxyfluorescein (6-FAM) labelled HH substrate was purchased at IDT and used without further purification. For the binding and activity assays, RNA was preheated (90 °C, 10 minutes), cooled down on ice, and added directly to the reaction.

### Lipid vesicle preparation

Lipid stocks dissolved in chloroform were pipetted into a glass vial and briefly evaporated under a steady flow of nitrogen gas. To remove organic solvent residues, a lipid film was dried under vacuum overnight. To obtain multi-layer vesicles (MLVs), lipid films were hydrated in the reaction buffer (10 mM HEPES pH 7, 5 mM CaCl_2_ and 5 mM MgCl_2_). Liposomes were shaken above the Tm of the lipids for one hour with brief vortexing every 15 minutes. A cloudy liposome suspension was transferred to the Eppendorf SafeLock tube and freeze-and-thawed 10x in liquid nitrogen/60 °C thermoshaker (1 minute/5 minutes, respectively), to reduce multi-lamellarity of the vesicles. Liposome suspension was then extruded 17x through 100 nm polycarbonate filters (Whatman) to achieve a consistent size distribution of vesicles. Liposome stocks were kept at 4 °C.

### Activity assays

In this study, we used lipid gel membranes (20:0 PC, DPPC; Tm = 66 °C and 41 °C, respectively), liquid disordered membranes (DOPC, DOPC:Chol 6:4), and lipid ordered membranes (DPPC:Chol 6:4) to investigate the effect of the lipids on the ribozyme activity and integrity. All of the binding and activity assays were performed in 10 mM HEPES pH 7, 5 mM CaCl_2_ and 5 mM MgCl_2_ unless stated differently (the presence of divalent ions is necessary to facilitate RNA-lipid interaction as well as RNA catalytic activity)^18,20–22,30,31,6^. HH (67 nM ribozyme, 133 nM substrate, 30 μL reaction) and HDV (14 nM, 30 μL reaction) incubations were conducted under temperature cycling conditions (10 cycles 60 °C -> 24 °C, 3 minutes each) to enhance RNA refolding^5^ (i.e. disrupt the non-specific base pairing between the ribozyme and the cleaved RNAs).

The overnight incubation of HDV-tRNA was performed under temperature cycling conditions (150 cycles) in the presence of 5 mM 20:0 PC (lipid gel membranes).

HH kinetic analysis in diluted conditions (5 nM ribozyme, 10 nM substrate) was performed with 100 μM lipids (either DPPC or 20:0 PC). The reactions were stopped after 3 temperature cycles (2 minutes 60 °C -> 8 minutes 24 °C). HH kinetic analysis in the reduced divalent ion content was performed in 10 mM HEPES pH 7, 500 μM MgCl_2_ and 500 μM CaCl_2_ in the presence of 250 μM lipids. The reaction yield was analysed after 1 cycle (2 minutes 60 °C -> 8 minutes 24 °C).

After the incubations, activity assays of HH and HDV were ethanol precipitated and analyzed using PAGE. HH activity was measured using the normalized product (P) intensity to sum of product and substrate (S) intensities:

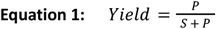

HDV activity readouts were based on the SybrGold-post stained gels using the ratio of un-reacted HDV-tRNA (Ht) band to the sum of tRNA (tRNA) and HDV-tRNA bands:

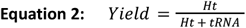

To calculate the relative activity in the presence of lipid vesicles, we quantified how much of the reaction substrate was processed in comparison with lipid-free samples. For the HDV-tRNA system the activity was fitted using the Hill’s equation to calculate K value:

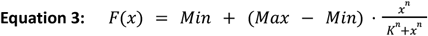

Min:minimal value

Max: maximal value

n: steepness of the fit

K: lipid-to-nucleotide ratio in which the relative reaction yield halved

### Binding assays

RNA was mixed with lipid vesicles and incubated at room temperature for 30 minutes. Samples were then centrifuged (128000 xG, 24 °C, 30 minutes) to separate lipid vesicles (pellet) from the supernatant. The supernatant was measured using miRNA Qubit quantification kit (Thermo Scientific). Obtained values were then normalized to the lipid-free sample, which represents the fraction of the RNA which is not bound to the vesicles. The binding assay result, binding efficiency, was expressed as a complement (1 minus the normalized values). Lastly, the measured binding values were fit to Hill’s equation in order to obtain K value (+/-SEM), which represents lipid-to-nucleotide ratio in which half of the RNA species are bound to the lipid vesicles.

For the binding and degradation analysis of HDV-tRNA, supernatant and pellet samples were analysed using PAGE, post-stained with SybrGold (activity analysis), and crystal violet (0.0025% crystal violet, 0.0005% Orange G in 10% ethanol staining, modified method from Yang et al.^32^, binding analysis). To estimate overall RNA binding we measured the lane intensities of supernatant (ISN) and pellet (IP) and determined the lipid:buffer partition coefficient (i.e. the preference of RNA to co-localise with lipid vesicles) values as follows:

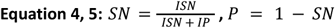

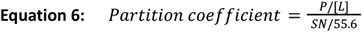

in which [L] is the half of the used lipid concentration (we assume that RNA interacts only with the outer lipid bilayer leaflet)^33,18^.

### Degradation readouts

RNA was incubated with different additives (liposomes, parts of lipid headgroup, lead acetate); after the incubation RNA was ethanol precipitated, separated on denaturing PAGE, and post-stained with SybrGold. Degradation was calculated as a ratio of degradation band intensities (Degr) and the sum of the the initial RNA (Ini) band and degradation band intensities:

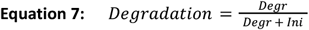

## Results

### The catalytic activity of RNA is modified by the presence of lipid membranes

We previously demonstrated an RNA-sequence dependent modification of ribozyme catalytic activity based on the presence of lipid gel membranes^18^. The model system was based on a trans-acting single-turnover R3C ligase and the activity changes were based rather on the substrate-lipid membrane interaction and not direct ribozyme-membrane interactions. Here, to diversify the potential modes of action, we chose different RNA with different mechanisms of activity: trans and cis acting ribozymes which catalyze trans- and cis-cleavage of RNA substrates, HH and HDV ribozymes, respectively. Both ribozymes bind to the lipid gel membranes with relatively high partition coefficient values (Equation 6, >1×10^6^)^18^ and with lipid:nt ratios around 1:1 (**Fig. 1a, b**).

**Fig. 1.**
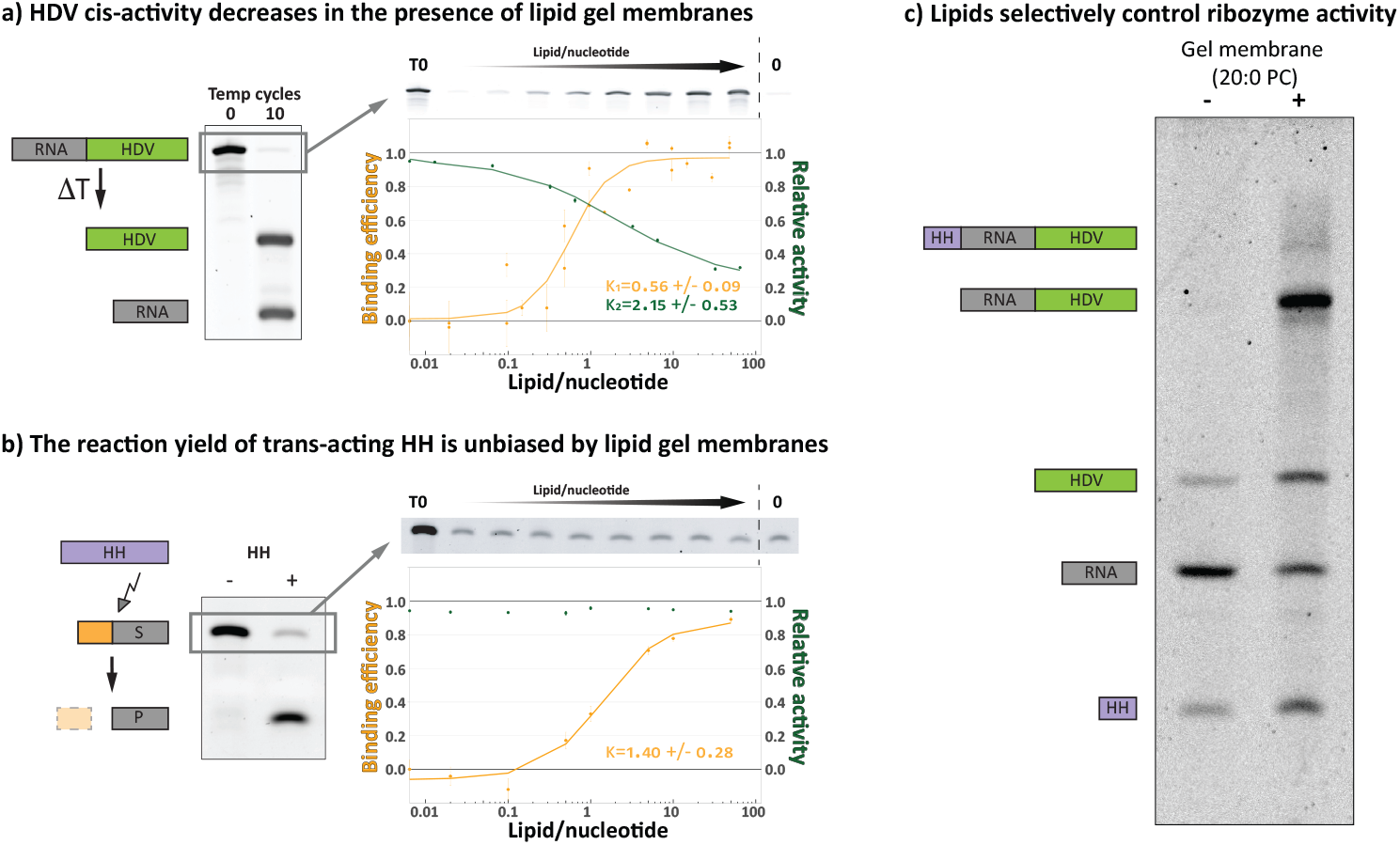
Lipid gel membranes act differently on different ribozymes. **a)** HDV ribozyme self cleaves under temperature cycling conditions (ΔT) into two products. The reaction yield (green) decreases with increasing lipid binding (either 20:0 PC or DPPC, orange). T0 represents the sample before the reaction, and 0 represents lipid-free incubation. All activity data points are the result of 10 temperature cycles. Binding and activity data, based on at least 3 assay replicates, were fitted to the Hill’s equation (solid line). Error bars represent SEM; please note that in case of activity assays (green points) errors were relatively small (<2% of the mean) thus they might appear underrepresented on the graph. **b)** HH ribozyme cleaves its substrate which generates two shorter RNA products. Despite significant lipid binding (DPPC, orange) there is no change in the total reaction yield (green). T0 represents the sample before the reaction, and 0 represents lipid-free incubation. All activity data points are the result of 10 temperature cycles. Binding data, based on 3 assay replicates, were fitted to the Hill’s equation (solid line). Error bars represent SEM; please note that in case of activity assays (green points) errors were relatively small (<2% of the mean) thus they might appear underrepresented on the graph. **c)** HH-tRNA-HDV fusion construct self-cleaves in two separate spots generating three products: HH, HDV, and tRNA. The presence of lipid gel membranes (20:0 PC) selectively and partially inhibits HDV activity generating half-cleaved intermediate (HDV-tRNA).

Activity assays of HDV and HH remarkably show different behaviors in the presence of lipid gel membranes. The HH final reaction yield was not affected, whereas the yield for the HDV ribozyme reaction decreased in a lipid concentration-dependent manner. Interestingly, the overnight incubation of the HDV construct with lipid gel membranes (20:0 PC) did not give a 100% reaction yield (**Suppl. Fig. 1**) which suggests that a fraction of HDV molecules bound to lipid gel membranes are unable to perform the cleavage reaction, most likely due to impaired refolding. Indeed, the HDV ribozyme can fold into multiple different structures (**Suppl. Fig. 2**) of which only a subset are catalytically active^5^. HH ribozyme, on the other hand, as a shorter molecule, folds into one dominant structure (**Suppl. Fig. 2**). The presence of lipid gel membranes most likely does not bias its folding, thus no differences in the final reaction yield were observed (**Fig. 1b**). These differential lipid-dependent responses open a potential opportunity to create a system in which RNA activity can be selectively influenced through interactions with lipid membranes. As a proof of principle, we used an HH-HDV construct^34^ in which both ribozyme types act together creating a characteristic digestion pattern. As expected, the presence of lipid gel membranes selectively inhibited the activity of HDV ribozyme whereas left HH functionality unaltered (reaction reached completion, **Fig. 1c**).

**Fig. 2.**
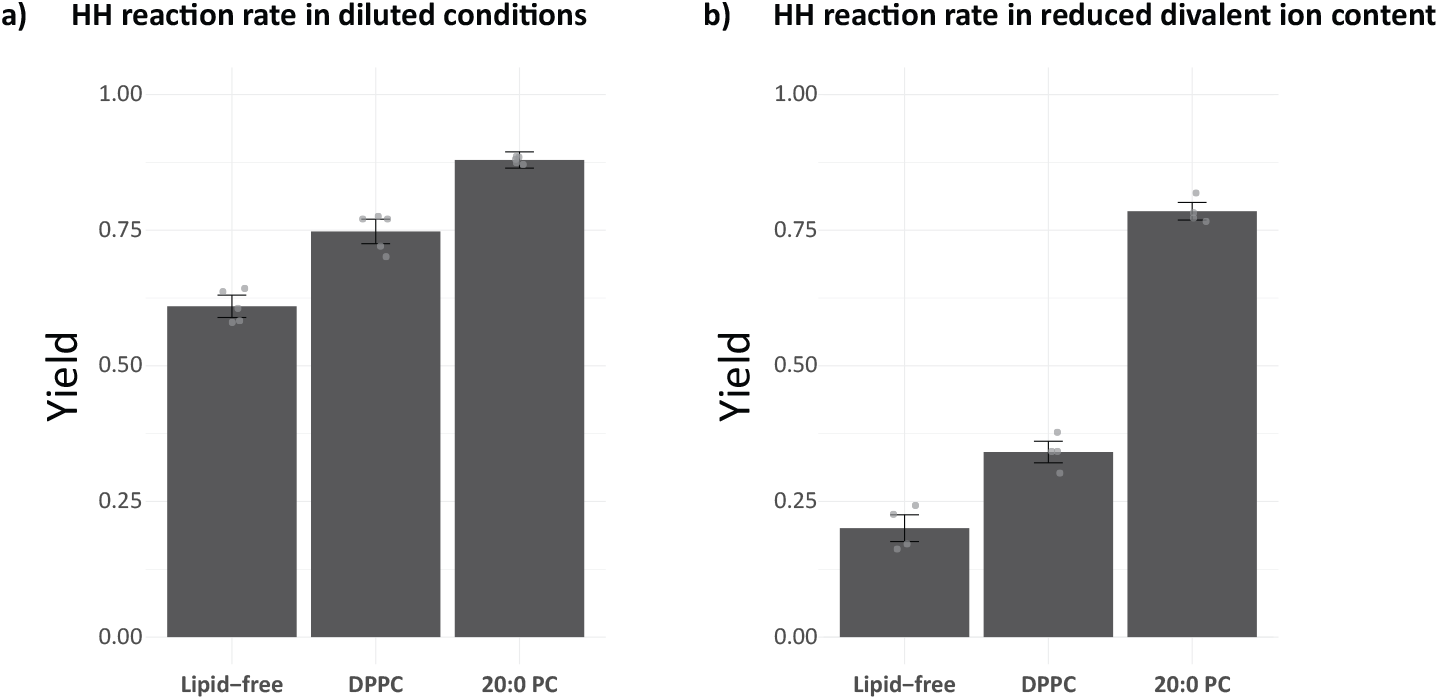
Lipid gel membranes increase the HH reaction rates. **a)** The HH ribozyme reaction in the diluted conditions (5 nM HH, 10 nM substrate) was stopped after 3 temperature cycles in which the reaction did not reach completion for the lipid-free system (**Suppl. Fig. 3a**). The reaction yield in this time point is significantly higher for the lipid-based systems. Obtained values are the average of 5 replicates (+/-SEM). **b)** The HH ribozyme in the lower divalent ion content (500 µM MgCl2, 500 µM CaCl2; 67 nM ribozyme, 133 nM substrate) was stopped after one temperature cycle in which the reaction did not reach completion for lipid-free system (**Suppl. Fig. 3b**). The presence of the lipid gel membranes yielded completion of the reaction (∼80%) whereas lipid-free reaction was still in progress (∼20%). Obtained values are the average of 4 replicates (+/-SEM).

In the above mentioned experiments we have determined that the presence of lipid gel membranes have no effect on the HH final yield (Fig. 1b, complete reaction after temperature cycling), however it is not clear if the presence of lipid gel membranes can affect the kinetics of the reaction. Thus, to determine if lipid gel membranes have any beneficial effect on HH ribozyme reaction rates (e.g. through up-concentration of RNAs on the lipid surfaces, increased RNA-RNA interactions^18,35^) we performed a time-lapse experiment. The assay was performed under diluted conditions (**Fig. 2a**), as well as with lower divalent ion concentration (**Fig. 2b**) to reduce the HH reaction rates. In both cases we observed that the overall reaction rates (yield over time, **Suppl. Fig. 3**) increased in the lipid gel membrane systems, namely 20:0 PC and DPPC. In this experiment, the gel membranes composed of DPPC exhibit a repeating lipid phase-transition from gel to liquid and back, since the reaction temperature was cycled above and below the lipid melting temperature (**Fig. 2, DPPC**). Thus, the reaction occurs in the presence of both liquid and gel membranes, the former of which more closely reflect the properties of fluid cell membranes. This demonstrates that the membrane does not necessarily have to be in a gel state over the entire course of the reaction; moreover, when the membrane is in a liquid phase, RNA can possibly diffuse off from the liquid membranes due to lower lipid binding affinity^18,22,36^ allowing the RNA to freely interact with other interaction partners not associated with lipid membranes.

**Fig. 3.**
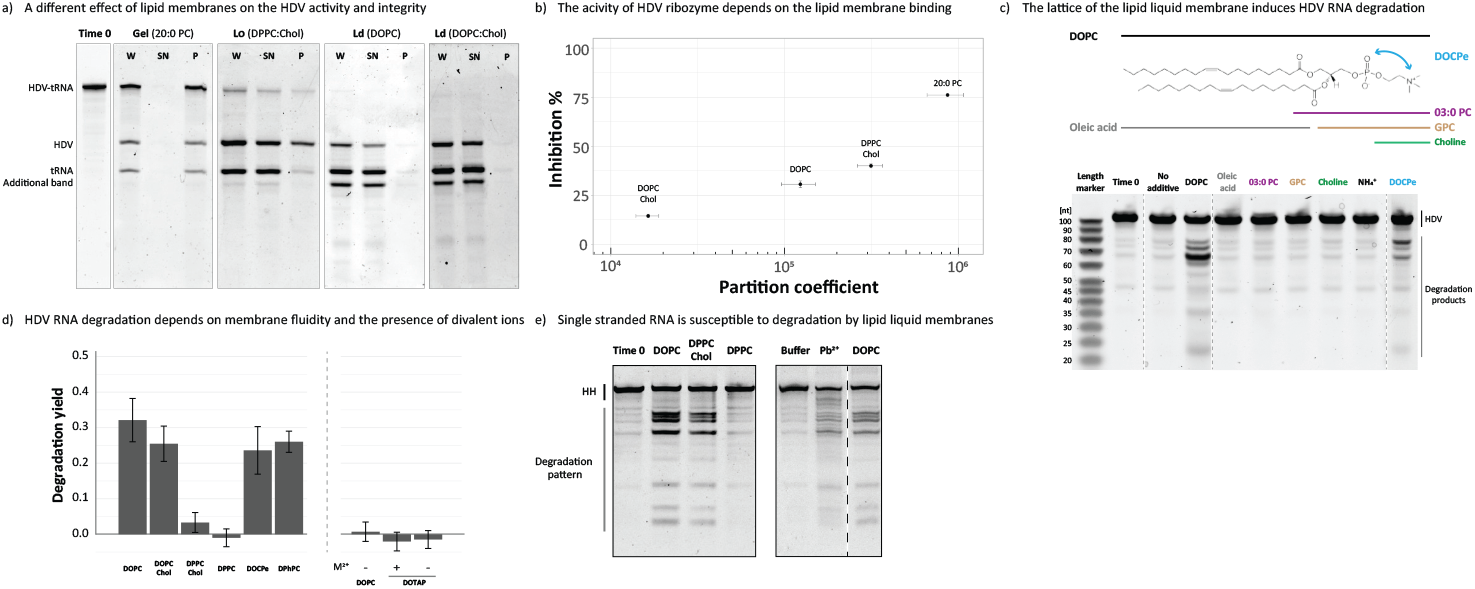
RNA activity and integrity is affected differently by different lipid membranes. **a)** HDV-tRNA construct (20 ng) was incubated in temperature cycling conditions in the presence of 1 mM lipids (gel – 20:0 PC, liquid disordered – DOPC, DOPC:Chol 6:4, liquid ordered – DPPC:Chol 6:4). RNA and lipids were then centrifuged and obtained fractions were analysed on the denaturing PAGE. W – reaction before separation, SN - supernatant (lipid free fraction), P – pellet (lipid bound fraction). Time 0 represents the sample before any activity and binding assay. **b)** The HDV inhibition was quantified and presented in the function of total RNA-lipid binding (partition coefficient). Lipid binding is correlated with the HDV inhibition with the largest effect for lipid gel membranes. Presented data is the average of 3 separate binding/activity assays (+/-SEM). **c)** 50 ng of HDV RNA was incubated in the presence of 1 mM additives at 24 °C for 24 hours. The presence of water-soluble additives (03:0 PC, GPC, choline, quaternary ammonium ion), as well as fatty acid-based aggregates (oleic acid) did not influence RNA integrity whereas liquid disordered lipid membranes (DOPC, DOCPe) induced RNA degradation. All assays were performed in the presence of 5 mM MgCl2 and 5 mM CaCl2. Time 0 stands for the RNA before the incubation. **d)** The HDV degradation was quantified and presented as a degradation yield. The presence of lipid liquid membranes (DOPC, DOPC:Chol, DOCPe, DPhPC) trigger RNA cleavage, whereas liquid ordered and gel membranes (DPPC:Chol, DPPC, respectively) do not enhance degradation. The presence of the metal divalent ions (M2+, namely MgCl2 and CaCl2), as well as PC-like headgroup geometry is crucial for RNA degradation: positively charged DOTAP does not induce RNA cleavage. Obtained data is an average of 3 separate assays (+/-SEM). **e)** 50 ng of HH RNA was incubated for 3 hours at room temperature with 1 mM lipids with different membrane fluidities. Liquid disordered (DOPC), liquid ordered (DPPC:Chol 6:4), gel phase (DPPC). Additionally, the RNA was incubated for 10 minutes in the presence of 10 mM lead acetate (Pb2+) which preferentially triggers RNA cleavage at the single-stranded parts of RNA. Dashed line on the right gel symbolizes that a middle lane was removed from the original gel image.

### The presence of lipid liquid membranes triggers RNA degradation

It was previously shown that most RNA sequences do not bind well to biologically relevant lipid liquid membranes^30,31,37,36,22,18^. However, Suga et al. reported that RNA binds to lipid liquid membranes and that this might affect RNA structure and catalytic activity^25,26,28^. Motivated by the unclear role of lipid liquid membranes in RNA-lipid systems we performed an activity assay in parallel with a binding assay of the HDV ribozyme. We hypothesized that liquid membranes should not affect RNA activity due to the lack of a stiff, RNA-structure-locking gel membrane. We observed high binding and inhibition of HDV activity for gel membranes (20:0 PC, **Fig. 3a**), which is in line with previous observations (**Fig. 1a**). Additionally, lower binding and smaller impact on HDV activity were observed in the presence of L_d_ membranes, while intermediate binding and inhibition of HDV were noted with stiffer L_o_ membranes (**Fig. 3a, b**). We noticed, however, that the HDV sequence undergoes degradation in the presence of L_d_ membranes generating a different RNA pattern than expected (**Fig. 3a, additional band**), and that degradation is dependent on the lipid concentration (**Suppl. Fig. 4a**). To investigate which part of the lipid membrane is responsible for the RNA degradation, HDV was incubated with different chemical components representing parts of the phospholipid (**Fig. 3c**). None of the components of the lipid headgroup can create stable lipid bilayers, and all are soluble in the buffer solution. The presence of lipid liquid membranes and divalent ions triggers RNA degradation: none of the lipid parts (fatty acids, parts of the lipid headgroup, **Fig. 3c**), and the incubation without divalent ions (**Fig. 3d**) affects RNA integrity. The RNA degradation is dependent on lipid membrane fluidity with highest degradation observed for lipid liquid membranes (DOPC, DOPC:Chol 6:4, **Fig. 3d**). Interestingly, the presence of liquid membrane-forming cationic lipid, DOTAP, in the absence and presence of divalent ions, does not induce RNA cleavage which suggests that a particular PC-like headgroup geometry and chemistry is the key for degradation. The incubation with the DOCPe lipid, which is an headgroup-inverted DOPC molecule, causes similar RNA degradation to DOPC. Lastly, the presence of the double bond is not necessary to induce degradation as incubation with a saturated DPhPC-based membrane in a liquid disordered state causes significant degradation (**Fig. 3d**). We noted that RNA degrades in a specific pattern (**Suppl. Fig. 4b**) rather than generating a smear of degradation products which most likely is due to structure-dependent RNA cleavage: typically single stranded parts of RNA are more prone to degradation^38,39^.

To investigate if the observed RNA degradation pattern represents selective cleavage of the single stranded segments we incubated the HH RNA (richer in single stranded parts compared with HDV) in the presence of lead acetate which preferentially induces the cleavage at the single stranded regions^40^; lead, as a divalent cation, binds to the RNA and, due to its charge distribution, induces a nucleophilic attack of 2’-OH ribose group on the adjacent phosphate causing hydrolysis.

We observed that lead-based digestion generated a pattern similar to lipid-dependent cleavage, however it also generated more degradation products (**Fig. 3e**). It is most likely that lead, as a metal ion, is smaller than lipid membranes and has better access to all parts of RNA compared with the bulky lipid membrane surfaces. We hypothesize that the presence of lipid liquid membranes allows single stranded parts of RNA to penetrate and transiently anchor the RNA in the proximity of the hydrophobic core of the membrane^24–28^ which changes the chemical environment (divalent ion distribution at the headgroup area, water activity) triggering specific RNA degradation, most likely utilising divalent ions bound to the lipid membrane. We observed different degradation patterns for DOPC and DOPC:Chol-based membranes which might be due to the different lipid headgroup geometry (**Suppl. Fig. 5**). Interestingly, we observed that Lo membranes, which caused negligible HDV degradation (**Fig. 3d, DPPC:Chol**), efficiently degraded HH (**Fig. 3e**), which is most likely due to the HH’s smaller size and higher content of single stranded parts (**Suppl. Fig. 2**) which can penetrate the L_o_ membrane. We have also observed different degradation patterns for DOPC and DOPC:Chol-based membranes which might be due to the different lipid headgroup geometry (Suppl. Fig. 5). Lastly, lipid gel membranes are harder to penetrate thus lower degradation rates could be due to inaccessibility of the hydrophobic core and deeper lipid headgroup areas.

## Discussion

RNA activity is involved in a diverse range of biologically relevant processes such as viral replication^2^, retrotransposon migration^4^, RNA maturation^1^, and gene expression control^17^. RNA activity has been shown to be regulated through the interaction with small solutes, as well as by proteins^9,11,15,17^. Here, we demonstrate the potential to control the activity of naturally occurring RNAs with lipid membranes.

We characterized how the activity of catalytic RNAs was affected by lipid membranes in a gel and liquid state, which significantly differ in fluidity. Our results extend previous findings that RNA-lipid interactions are sensitive to several factors such as membrane fluidity, lipid composition, and RNA sequence and structure^18,22,30,31,36,37^. In agreement with the published data^36,22,18^, we observed the highest binding for lipid gel membranes (partition coefficient ∼1e6), and lowest for liquid disordered membranes (partition coefficient ∼1e4 to ∼1e5, **Fig. 1, Fig. 3a, b**). Despite the higher lipid order we did not observe increased binding of RNA to DOPC:Chol based membranes; in contrast, binding to pure DOPC membranes was higher compared with DOPC:Chol membranes, which might be due to the the presence of a less regularly distributed DOPC molecules (i.e. dilution of DOPC with cholesterol molecules) or other structural effects. Interestingly, we observed the correlation of the lipid membrane binding with the inhibition of the cis-acting HDV ribozyme activity (**Fig. 3b**). The interaction with lipid membranes can influence RNA structure which could, in turn, also affect RNA activity. The lipid-based HDV activity inhibition is likely due to an effect on RNA refolding in the presence of a stiff gel phase lipid bilayer^41^; alternatively, steric hindrance from the bulky lipid membranes as well as direct nucleotide-lipid interactions may have played a role in the HDV inhibition^23,42–45^ as a slight effect was also observed for lipid liquid membranes (**Fig. 3a, b**).

HH remained active and even accelerated under similar conditions (**Fig. 1b, Fig. 2, Suppl. Fig. 3**). HH ribozyme activity involves the recognition of the substrate through base pairing with the ribozyme^46^. Lipid membranes, in this case, can act as a scaffold^18,35^ templating the HH’s substrate-recognition part, facilitating HH-substrate binding and, eventually, substrate cleavage. Increased HH activity in the presence of lipids might be also caused by the increased co-localisation of HH and its substrate on lipid membrane surfaces locally increasing the concentration of both reactants^47^, which might be a larger factor under more diluted conditions (**Fig. 2a**). Enhanced HH activity in the presence of lipid gel membranes and reduced divalent ion content (**Fig. 2b**), on the other hand, might be caused by the increased local divalent ion concentration and their favorable distribution (spacing between lipid membrane headgroups)^48^.

Modification of RNA activity by lipids has been reported before. For instance, the activity of L1 ligase was modified by the fatty acid based lipid aggregates^49,50^. Similarly, RNA polymerase ribozyme activity was tuned by artificial lipid binding^47^. Lastly, R3C ligase activity was modified by the presence of lipid gel membranes^18^. In those cases it has been speculated that the presence of lipids modify the RNA microenvironment (RNA crowding on lipid membrane surfaces, divalent ion binding), as well as increase the probability of RNA-RNA interactions in which lipid membranes act as a scaffolding platform for RNA species increasing the probability of base-pairing.

Remarkably, the presence of biologically relevant lipid liquid membranes affected RNA integrity, resulting in partial cleavage with characteristic patterns that depended on the presence of the lipid liquid membrane scaffolds (**Fig. 3**). Lipid based RNA degradation might be caused by several factors; it was shown that lipid membranes can bind metal ions, including divalent ions such as Mg and Ca^48,51^. It was also shown that RNA degradation is boosted in the presence of divalent ions^52^ as well as within single stranded regions^39^. Higher degradation rates for lipid-based systems might be due to the specific spacing of divalent ions allowing more efficient RNA cleavage. The cleavage in single stranded fragments might be due to the partial penetration of the headgroup area of the lipid membrane; indeed, it was shown that ssRNA can slightly penetrate the membrane^24,27,25,26^ which, together with specific metal ion distribution, might be the main reason for degradation.

Our results suggest that the viroid, HDV, and retrozyme replication could also potentially be affected by the presence of the lipid membranes. Our observations of simple RNA-lipid based systems create a framework to understand if and how lipid membranes can be involved in RNA metabolism. For instance, mRNA degradation might occur at specific membrane fluidities, either activating or inhibiting its translation activity. Additionally, lipid-sensitive riboswitches might represent another level of gene expression in which RNA is activated by lipid membranes with specific composition or property. Lipid-membrane co-localized RNAs have been found *in vivo* in the form of mRNAs^53–55^, lncRNAs^56,57^, and glycoRNAs^58^. Recently it was also shown that cell-extracted RNA species can bind to lipid membranes^59^. In some cases, the mechanism of binding is not entirely clear, however it was proposed that some of the RNA sequences can have affinity for lipid membranes^30,31,37,60^. The lipid-driven RNA-activity changes might have several implications in cellular biology. The close proximity of catalytic RNAs, such as ribosomes, to different lipid membranes might have an influence on RNA activity and integrity which could further influence cellular homeostasis. It was demonstrated that lipids forming the nuclear envelope increase RNA stability^61^ which, for example, might improve the activity of RNAseP, a ribozyme involved in the small RNA processing.

Lipid-based tuning of the RNA stability and activity could also have implications in primordial scenarios in which complex protein-based processes were not present. For example, templating and cleavage activities of the lipid membranes could act similarly to the RNA-induced silencing complex (RISC)^62^; lipid-bound RNA would be exposed to interactions with target RNAs which would trigger their degradation. in this way, such a system acts as a primitive lipid-based RNA interference system. Moreover, lipid-based RNA degradation could possibly act as a lipid membrane fluidity biosensor^63^ in which changes in lipid membrane fluidity and composition could facilitate RNA cleavage generating a pool of primitive secondary signalling molecules similar to protein-based signalling cascades. Additionally, the lipid membrane-based enhancement of HH catalytic activity could act as a primitive version of membrane associated RNAses, such as RNAseE^64^. Membrane recruited RNAseE is a part of RNA degradosome and lipid membranes stabilize its activity. Similar patterns could be observed in the RNA-lipid based system in which the membrane recruited HH ribozyme is more prone to degrade its substrates (**Fig. 2**) which could act as a primordial RNA regulation mechanism (temperature cycling in this case could mimic the presence of thermal vents in the prebiotic scenarios). Lastly, the inhibition of HDV ribozyme and activation of HH by lipid gel membranes resembles allosteric regulation processes (for instance non-competitive inhibition) that are part of the enzyme-based reactions.

Development of lipid-binding RNAs could potentially have applications for the artificial introduction of lipid-binding RNA species *in vivo*. For example, the mRNA translation process could be co-localized with the lipid membranes providing a means to control the localization of membrane protein translation^65,54^. Creating new aptazymes and bi-functional aptamers could also create diverse and controllable functional outcomes; it was reported that merging the lipid-binding sequences with tryptophan aptamers increased tryptophan lipid membrane permeability^66^. Modifying existing RNAs with lipid-binding motifs might play a role in the exosome packing processes^60,67–69^. Novel lipid-based aptazymes and riboswitches could generate a pool of lipid-dependent feedback loops controlling further the stability and activity of modified ribozymes and mRNAs. Lastly, a bioinformatic analysis of lipid-binding motifs in currently developed lipid-binding RNAs would help to find naturally existing lipid-binding RNAs.

Merging our observations of modified RNA activity (R3C ligase, HH and HDV ribozymes) and altered RNA stability (including RNA degradation and lipid-based cleavage of other RNA species) in the presence of lipid membranes provides a framework for developing functional systems based on the interactions between RNA and lipids. This framework lays the groundwork for an understanding of RNA-lipid interplay in living cells, as well as in synthetic and prebiotic systems.

## Acknowledgements

The authors would like to thank 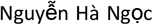 Anh for the constructive feedback, and Mario Mörl lab for helpful discussions, as well as for the aliquot of the HH-tRNA-HDV construct. This work was supported by the B CUBE of the TU Dresden, a German Federal Ministry of Education and Research BMBF grant (to J.S., project 03Z22EN12), and a VW Foundation ‘‘Life’’ grant (to J.S., project 93090).

## Author contributions

T.C. performed and analyzed experiments, conceptualized and wrote the manuscript.

J.P.S acquired the funding, conceptualized, and edited the manuscript.

## Conflict of interest

The authors declare no conflict of the interest.

## Notes

### Competing Interest Statement

The authors have declared no competing interest.

### Summary of Updates

Revised manuscript text. Minor edits, and some additional text in the intro, results, and discussion.

