## Supplementary Figures for "Effects of lipid membranes on RNA catalytic activity and stability"

**Supplementary Table 1:** T7 transcription templates (5’ -> 3’) and buffer components (**bold**: T7 promoter)

| | |
|---|---|
| Schistosoma hammerhead DNA template for T7 transcription | GGAGGGCATTTCGTCCTATTTGGGACTCGTCAGCTGGATGTACCTCGC**CTATAGTGAGTCGTATTAG** |
| Sequence of HH-tRNA-HDV DNA template | **TAATACGACTCACTATAG**GGAGAAATCCGCCTGATGAGTCCGTGAGGACGAAACGGTACCCGGTACCGTCGCGGATTTAGCTCAGTTGGGAGAGCGCCAGACTGAAGATCTGGAGGTCCTGTGTTCGATCCACAGAATTCGCAGGGTCGGCATGGCATCTCCACCTCCTCGCGGTCCGACCTGGGCTACTTCGGTAGGCTAAGGGAGAAGCTTGGCACTGGCCGTCGTTTAAGGGCGAATTCTGCAGAT |
| Substrate of Schistosoma hammerhead | 6-FAM-GGAGGGCAUCCUGGAUUCCACUCGCC |
| T7 transcription buffer | 40 mM Tris pH 8<br>23.5 mM MgCl2<br>15 mM DTT<br>10 mM NaCl<br>2 mM spermidine<br>7.5 mM nucleotides (1.88 mM each)<br>5 U/ml inorganic pyrophosphatase (NEB, M0361)<br>0.018 g/L T7 polymerase (homemade, MPI-CBG) |

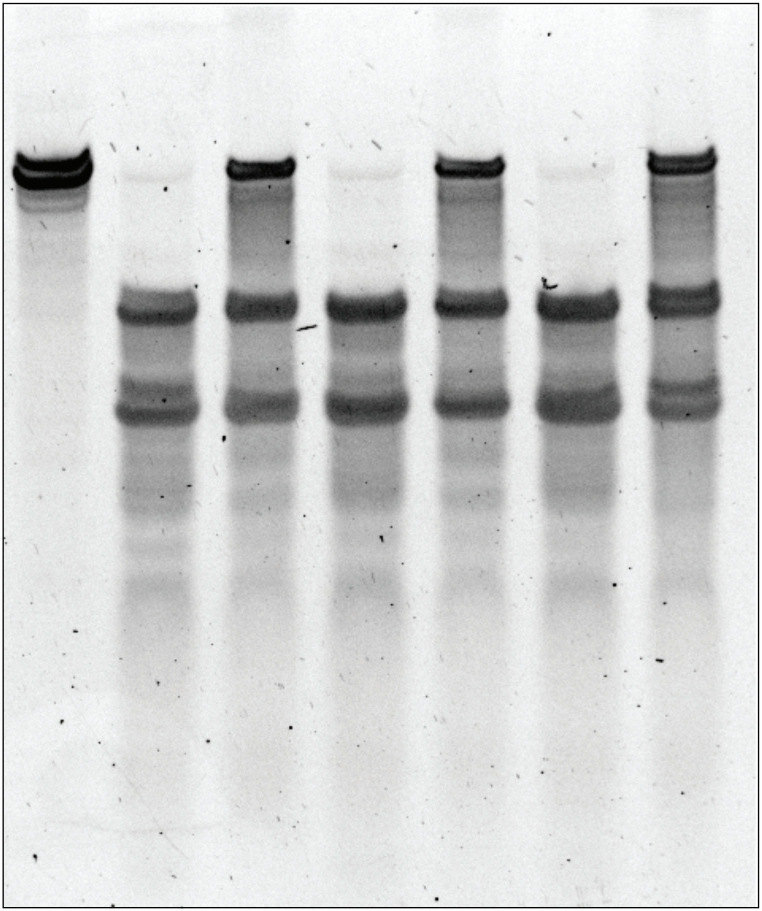


**Supplementary Figure 1 HDV ribozyme is partially inhibited by lipid gel membranes after overnight temperature cycling (150 cycles).** The reaction was performed in the buffer with 5 mM 20:0 PC (gel membranes), reaction triplicates are presented.

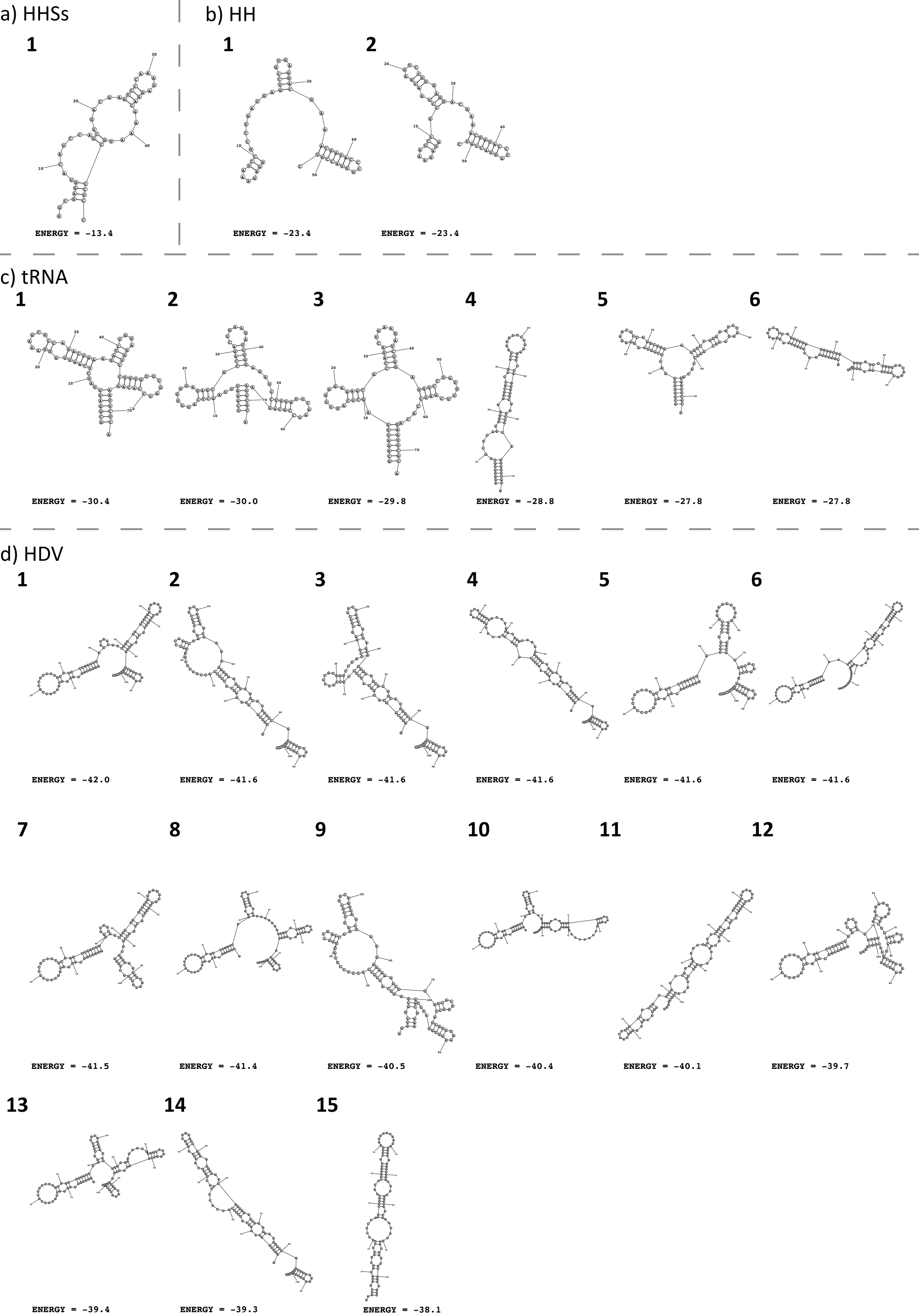


**Supplementary Figure 2 Secondary structures of used RNAs generated in silico using fold tool[70].**
**a)** Schistosoma hammerhead (HHSs) used as trans-acting ribozyme generates one dominant structure. **b)** Hammerhead generates similar two structures. **c)** tRNA generates multiple (6) different structures, dominantly double stranded. **d)** HDV generates the largest number of different structures (15), mostly double stranded. HH, tRNA, and HDV are part of the HH-tRNA-HDV construct. Presented energy values are free energy of structures in kcal/mol.

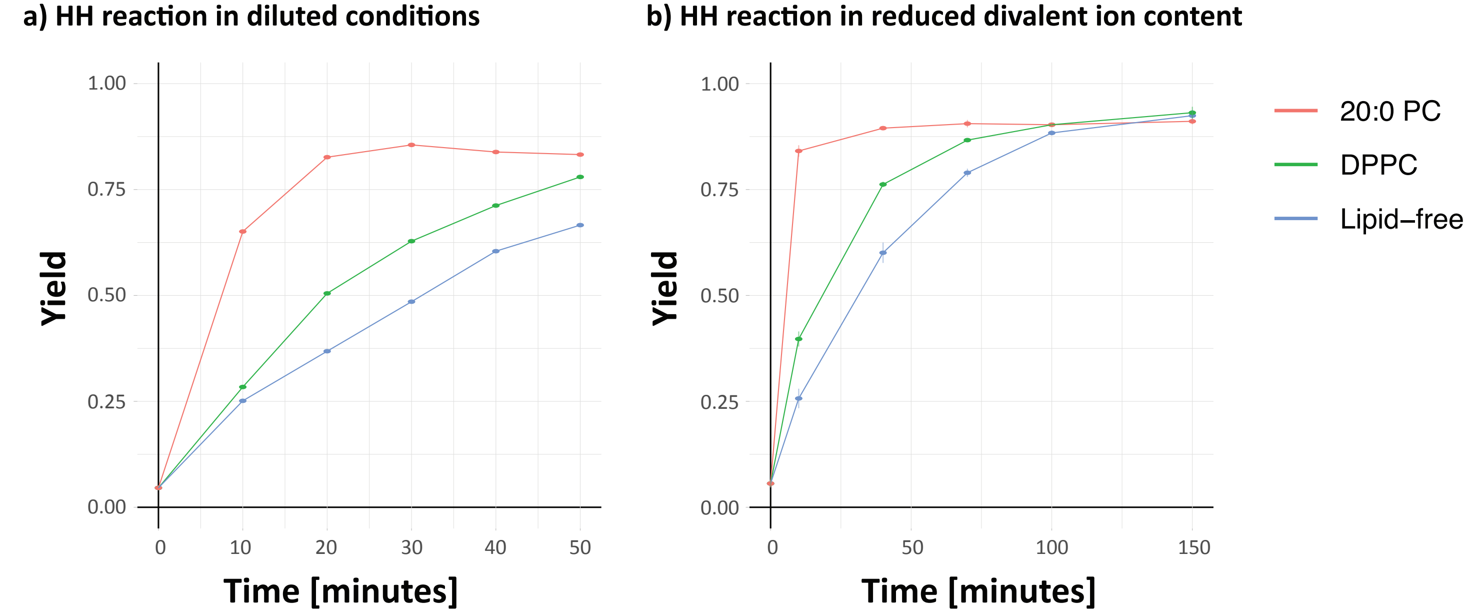


**Supplementary Figure 3 HH ribozyme reaction rate increases in the presence of lipid membranes. a)** Test experiment (n=1) of HH reaction in the diluted conditions (5 nM ribozyme, 10 nM substrate) shows that the presence of lipid gel membranes improves reaction rate; the largest effect was visible after 20-40 minutes. This experiment was performed once to determine the time point in which the differences of the HH reaction yield were visible for the lipid and lipid-free system, thus lacking error bars. **b)** HH reaction rate in the decreased divalent ion content (500 µM $MgCl_2$, 500 µM $CaCl_2$) is faster in the presence of lipid gel membranes and the largest differences are visible after 10-40 minutes of the temperature cycling (1-4 cycles). Error bars are SEM from 4 replicates and are relatively small (<3%), which might bias their readout.

**a) Degradation of HDV ribozyme in the presence of lipid liquid membranes**

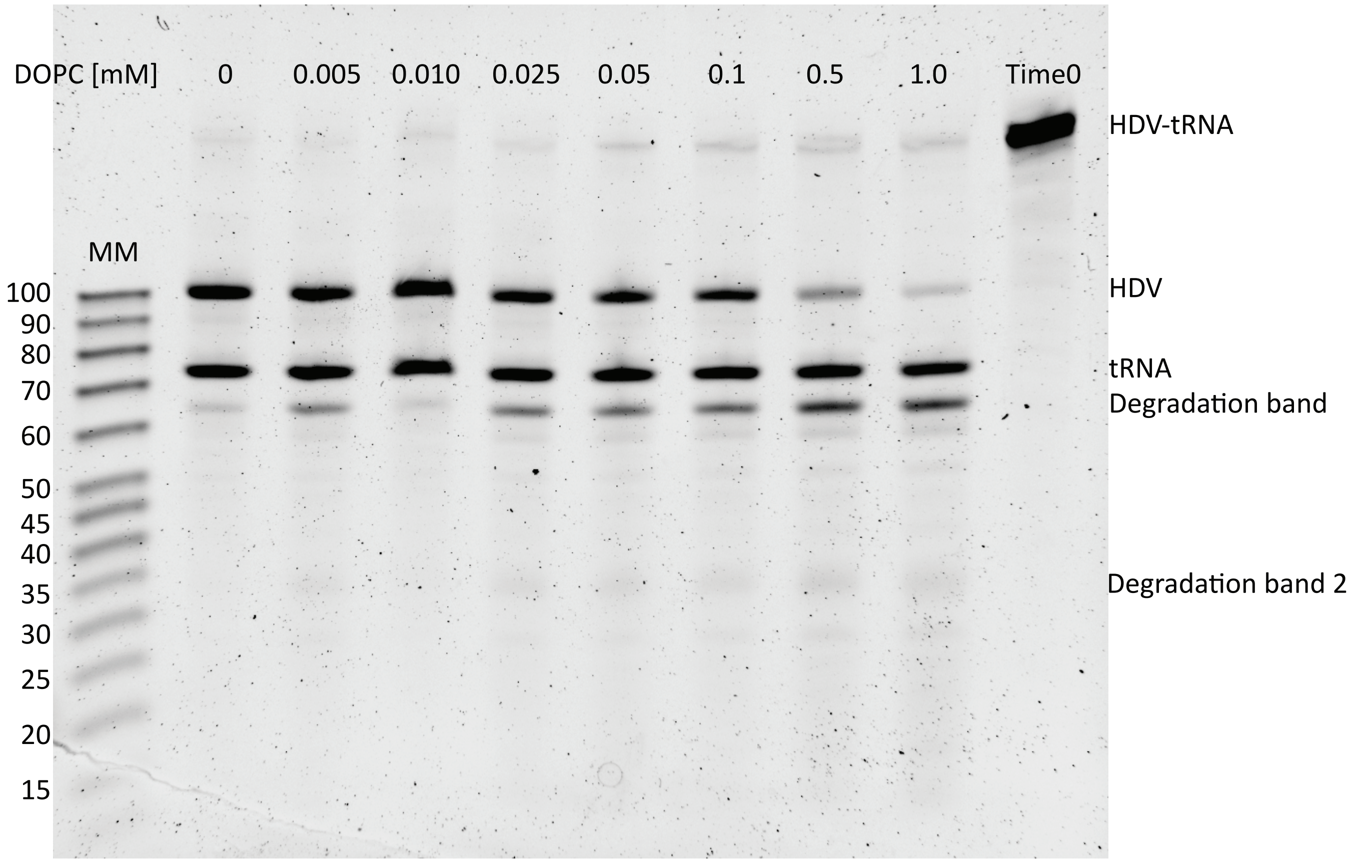


**b) Length estimation of the degradation products**

| length [nt] | log length | migration [pixels] |
|---|---|---|
| 100 | 2 | 509 |
| 90 | 1,95424251 | 616 |
| 80 | 1,90308999 | 738 |
| 70 | 1,84509804 | 884 |
| 60 | 1,77815125 | 1058 |
| 50 | 1,69897 | 1265 |
| 45 | 1,65321251 | 1381 |
| 40 | 1,60205999 | 1520 |
| 35 | 1,54406804 | 1671 |
| 30 | 1,47712125 | 1834 |
| 25 | 1,39794001 | 2016 |
| 20 | 1,30103 | 2234 |

| RNA | migration [pixels] | Approx. Length |
|---|---|---|
| HDV | 506 | **99,9175894** |
| tRNA | 833 | **74,0541924** |
| Degradation1 | 982 | **64,6058909** |
| HDV | 512 | **99,3699184** |
| tRNA | 837 | **73,7833393** |
| Degradation1 | 984 | **64,4876347** |
| Degradation2 | 1715 | **33,0109861** |

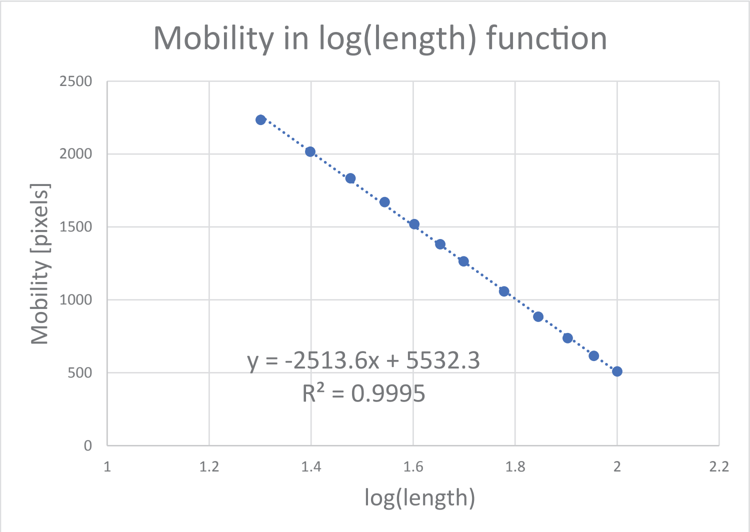


**Supplementary Figure 4 HDV ribozyme degrades in the presence of lipid liquid membranes in a lipid concentration dependent manner. a)** 20 ng of HDV-tRNA construct was incubated with various concentrations of DOPC lipid membranes (liquid lipid membranes) in temperature cycling conditions. RNA was then analysed using PAGE (run along with MM - length marker, Affimetrix, Ref: J76410, Lot: 4310399), presented lengths are in nucleotides. **b)** The marker migration was quantified as pixels in the whole lane, and the plot of migration (pixels) in the function of log(length) was generated ($R^2 = 0.9995$). Using the migration values for different bands and using a linear fit of the marker bands (graph) the lengths of the RNA bands presented on the gel were calculated. The degradation band 1 has a length ~ 65 nucleotides, whereas degradation band 2 ~ 35 nucleotides, which sums up to HDV length ~ 100 nucleotides.

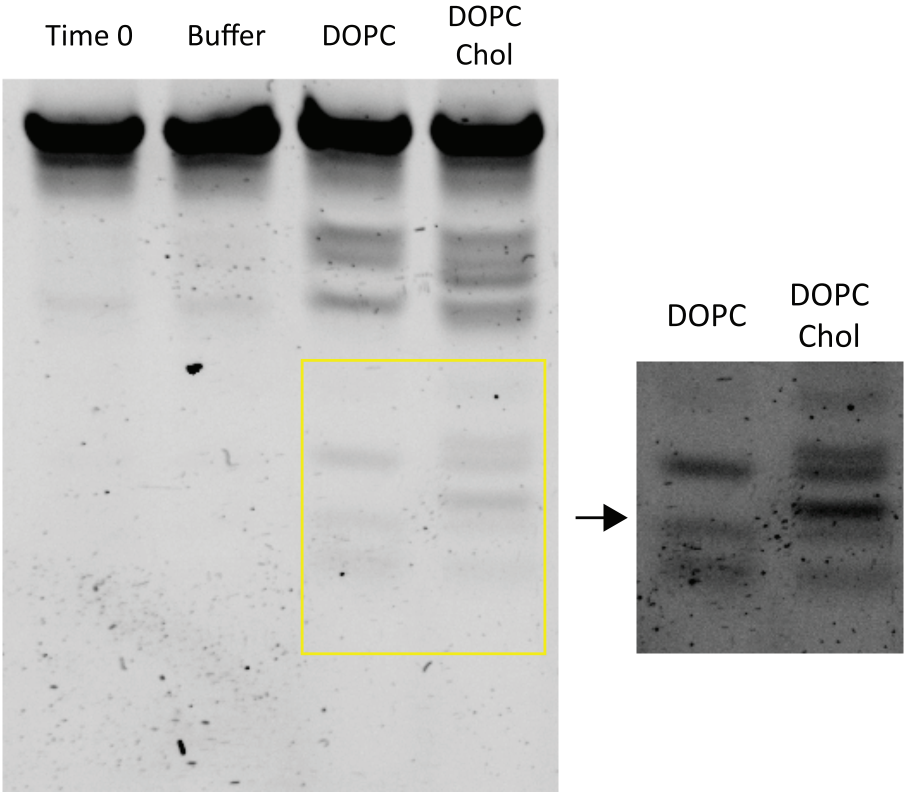


**Supplementary Figure 5 The HH degradation pattern is different for different liquid membranes.** HH was incubated with DOPC and DOPC:Chol membranes: both lipid membranes degrade HH, however the generated degradation pattern differs.
